# The *Drosophila* G protein-coupled receptor, GulpR, is essential for lipid mobilization in response to nutrient-limitation

**DOI:** 10.1101/2025.04.19.649675

**Authors:** Daniela Barraza, Xiang Ding, Zihuan Wang, Bat-Erdene Jugder, Paula I. Watnick

## Abstract

Enteroendocrine cells (EECs) of the intestinal epithelium are major regulators of metabolism and energy homeostasis. This is mainly due to their expression and secretion of enteroendocrine peptides (EEPs). These peptides serve as hormones that control many aspects of metabolic homeostasis including feeding behavior, intestinal contractions, and utilization of energy stores. Regulation of EEP production and release depends largely on EEC-exclusive G protein-coupled receptors (GPCRs) that sense nutrient levels. Here we report the identification of a GPCR expressed principally in EECs, which we have named GulpR due to its role in the response to nutrient stress. We show that GulpR regulates transcription of the EEP Tachykinin (Tk) and that both GulpR and Tk are essential for the transcriptional response that promotes survival of nutrient limitation. Infection with *V. cholerae* also activates transcription of Tk and lipid mobilization genes. While GulpR is required for activation of Tk transcription during infection, Tk does not play a role in regulation of lipid mobilization genes or survival of infection. Our findings identify a role for GulpR and Tk in survival of starvation and suggest that, although starvation and infection both require significant mobilization of energy stores, the signal transduction systems that regulate the metabolic response to each are distinct.

**Author Summary.:** Humans and other animals, including *Drosophila*, metabolize dietary nutrients such as sugars, lipids, and proteins into polysaccharides, fatty acids, and amino acids, respectively, to generate energy that fuels essential cellular processes like cell division, ion transport, muscle contraction, and more. The ability to adapt to changes in nutrient availability and energy demand is therefore crucial for homeostasis and survival. Nutrient scarcity during starvation and an increased demand for energy during an immune response against pathogenic infection require utilization of the body’s own lipid and glycogen stores. This adaptive response largely relies on the ability of the intestine to sense and respond to a variety of stimuli, including microbes and dietary nutrients. Here, we have identified and characterized a *Drosophila melanogaster* receptor that is expressed in a rare intestinal cell type. We report that this receptor regulates production of peptide hormones that are known to impact metabolic homeostasis and discover that one of these peptide hormones is crucial for utilization of systemic lipid stores when flies experience starvation but not infection stress. Our findings therefore indicate that activation of lipid mobilization in response to nutrient limitation and infection are regulated via different mechanisms.

## Introduction

Maintenance of metabolic homeostasis is crucial for the health span and life span of living organisms. This requires matching food consumption and catabolism to the energy expended. This balance is achieved in part via the action of gut-derived peptide hormones produced in enteroendocrine cells (EECs) [1]. EECs comprise the largest endocrine organ in the body, secreting more than 20 different enteroendocrine peptides (EEPs) [2–4]. EEPs can act in a paracrine fashion to regulate nutrient absorption, gut motility, epithelial renewal and other processes within the intestinal epithelium or in an endocrine fashion to regulate appetite, satiety, metabolism, and energy expenditure. Appropriate EEP expression and secretion relies on the ability of EECs to sense nutritional status and nutrient intake via systemic and intestinal signals, respectively. This is achieved, in large part, via the action of cell-autonomous G protein-coupled receptors (GPCRs). GPCRs are cell membrane-associated receptors that activate a signal transduction cascade in response to binding small molecules or peptides. In EECs, GPCRs respond to nutrients, metabolites secreted by the intestinal microbiota, bile acids, neuropeptides, or other molecules by regulating EEP expression and exocytosis-mediated secretion [3, 5, 6].

*Drosophila melanogaster* has been used extensively as a model organism for the study of gastrointestinal (GI) processes as its intestine shares structural and functional homology with the mammalian GI tract [7, 8]. The adult *Drosophila* intestinal epithelium is subdivided into foregut, midgut, and hindgut sections [9]. The *Drosophila* midgut epithelium is composed of three mature cell types: Intestinal stem cells, enterocytes (ECs), and enteroendocrine cells (EECs). While ECs are most abundant, EECs account for 5-10% of cells in the intestinal epithelium [9]. As in humans, *Drosophila* EEPs are important regulators of intestinal and systemic physiology and metabolism [10–12].

The *Drosophila* midgut can be subdivided into three regional sections (anterior, middle, posterior), exhibiting unique functional properties [9, 13]. The microbiota of the fly resides mainly in the anterior midgut (AMG), which is also the compartment responsible for metabolism of complex macromolecules such as sugars, fats, and proteins. Here we focus on regulation of the EEP Tachykinin (Tk), which is essential for innate immune signaling and lipid homeostasis in the AMG [10, 14, 15]. Tk has also been shown to impact lipid stores in the *Drosophila* fat body, which is the main site of energy storage in the fly and functionally analogous to human adipose tissue and liver [10, 16–18].

To survive starvation or infection, animals must mobilize stores to meet their energy requirements [16–18]. Here we describe an orphan GPCR we have named GulpR that is specifically expressed in EECs, regulates Tk expression in response to nutrient stress and infection, and is essential for survival of starvation but not infection. These findings suggest that, while both the response to starvation and infection require considerable energy expenditure, distinct signaling pathways participate in generating the metabolic response to each. Thus, the type of energetic demand encountered determines the signaling pathway activated.

## Results

### A GulpR mutant displays decreased numbers of Tk-expressing (Tk+) cells in the intestine and increased lipid storage in the anterior midgut

In a previous RNA-seq experiment using *Vibrio cholerae*-infected whole flies, we noted a positive correlation between the intestinal levels of the short-chain fatty acid (SCFA) acetate during infection and expression of an EEC-specific orphan GPCR CG32547, which we have named GulpR (CG32547) [19, 20] (Fig 1A). GulpR has two predicted transcripts both consisting of 8 exons and 7 introns but with differing 3’ untranslated regions. These encode identical polypeptides. Because acetate prevents lipid accumulation in the *Drosophila* AMG by increasing Tk expression, we questioned whether GulpR might be involved in regulating Tk expression and/or release [14]. To test this hypothesis, we first characterized an available *GulpR* transposon-insertion mutant fly (*GulpR*^f06408^). This transposon is located at the beginning of intron 5 and, therefore, interrupts both transcripts. We began by confirming that *GulpR* transcription was reduced in this mutant as compared to control *y^1^w^1^ (yw*) flies (Fig 1B). We then questioned whether *GulpR* was expressed in Tk+ EECs. To test this, we measured transcription of *GulpR* in the intestines of Tk> control and Tk>*GulpR*^RNAi^ flies. In fact, compared with Tk> control flies, *GulpR* expression was decreased by approximately 40% (Fig 1C). This suggests that *GulpR* is present in Tk+ cells and likely other EECs. We then quantified Tk+ EECs and lipid accumulation in the intestines of control *yw* and *GulpR^f06408^* mutant flies. We observed an accumulation of lipids in the AMG of the *GulpR* mutant but not the PMG. This was accompanied by a reduction in the number of Tk+ EECs in both the AMG and PMG (Fig 1D-F). We conclude that GulpR regulates Tk expression and/or secretion in both the AMG and PMG, but lipid accumulation only in the AMG.

**Figure 1:**
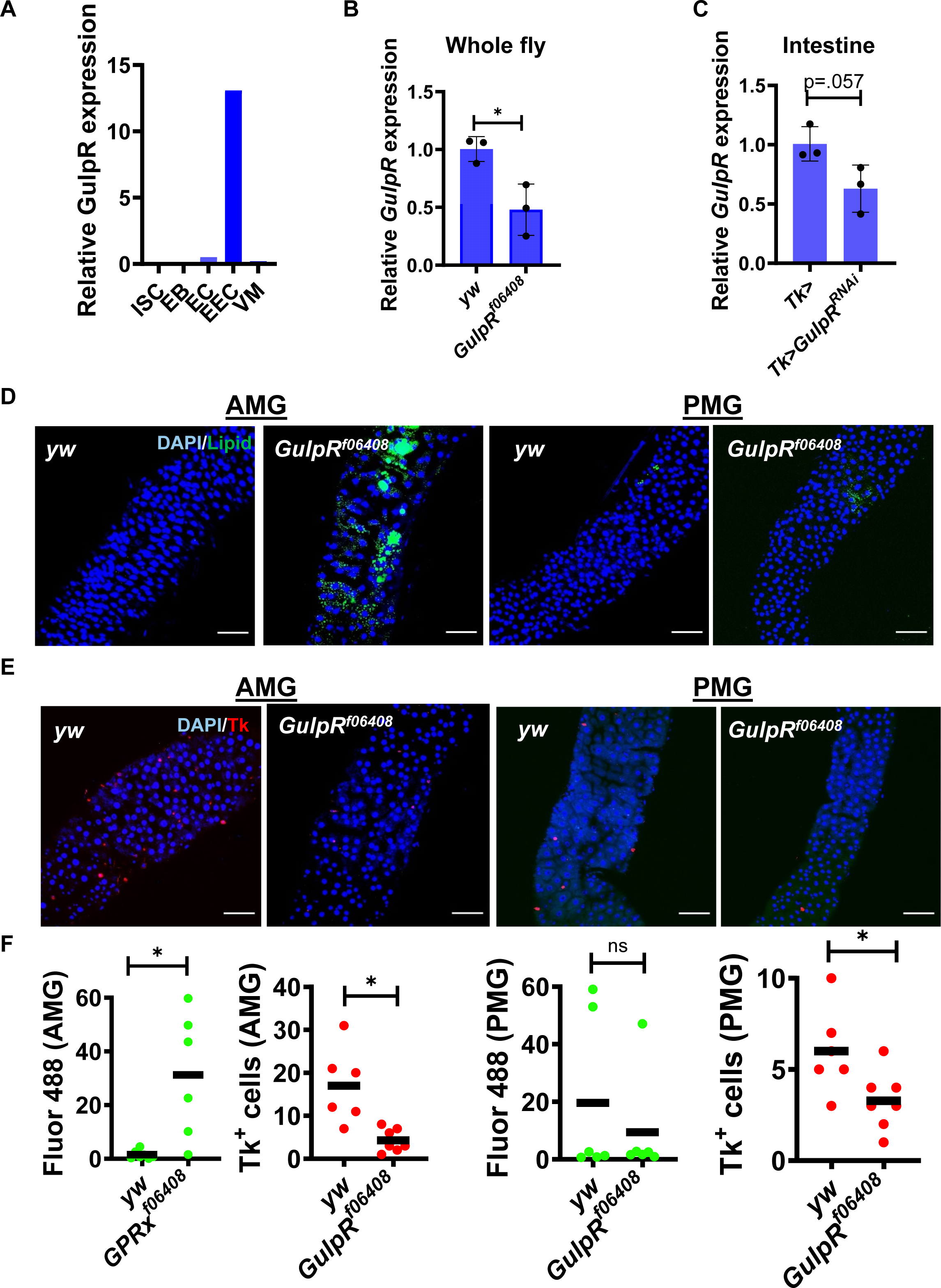
A *GulpR* mutant displays increased intestinal lipid storage and decreased Tk+ EECs. (A) GulpR expression in various cells of the midgut (from http://flygutseq.buchonlab.com/). (B) qRT-PCR analysis of *GulpR* transcription in whole control *yw* or *GulpR*^r06408^ flies. (C) qRT-PCR analysis of *GulpR* transcription in the intestines of control Tk> or Tk>*GulpR*^RNAi^ flies. The mean of biological triplicates is shown. Error bars represent the standard deviation. A student’s t test was used to assess significance. (D) Representative fluorescence images showing DAPI and BODIPY(Lipid) staining in the anterior midgut (AMG) and posterior midgut (PMG) of the indicated fly genotypes. Scale bars, 50 μM. (E) Representative immunofluorescence images showing DAPI and TK staining in the anterior midgut (AMG) and posterior midgut (PMG) of the indicated fly genotypes. Scale bars, 50 μM. (F) Quantification of total fluorescence and Tk+ EECs in the AMG and PMG of the indicated fly genotypes. Female flies were imaged. The mean of at least six intestines is shown. A Welch’s t test was used to assess significance. * p < 0.05, ns not significant.

### *GulpR*^RNAi^ in Tk+ EECs decreases intestinal Tk expression and increases lipid accumulation

To determine whether cell-autonomous GulpR expression is required to maintain Tk levels and lipid homeostasis in the *Drosophila* intestine, we compared the phenotype of Tk> control flies with Tk>*GulpR*^RNAi^ flies. Like the *GulpR* mutant, Tk>*GulpR^RNAi^* flies exhibited an increase in lipid storage in the AMG and a decrease in the number of Tk+ EECs in the AMG and PMG (Fig 2).

**Figure 2:**
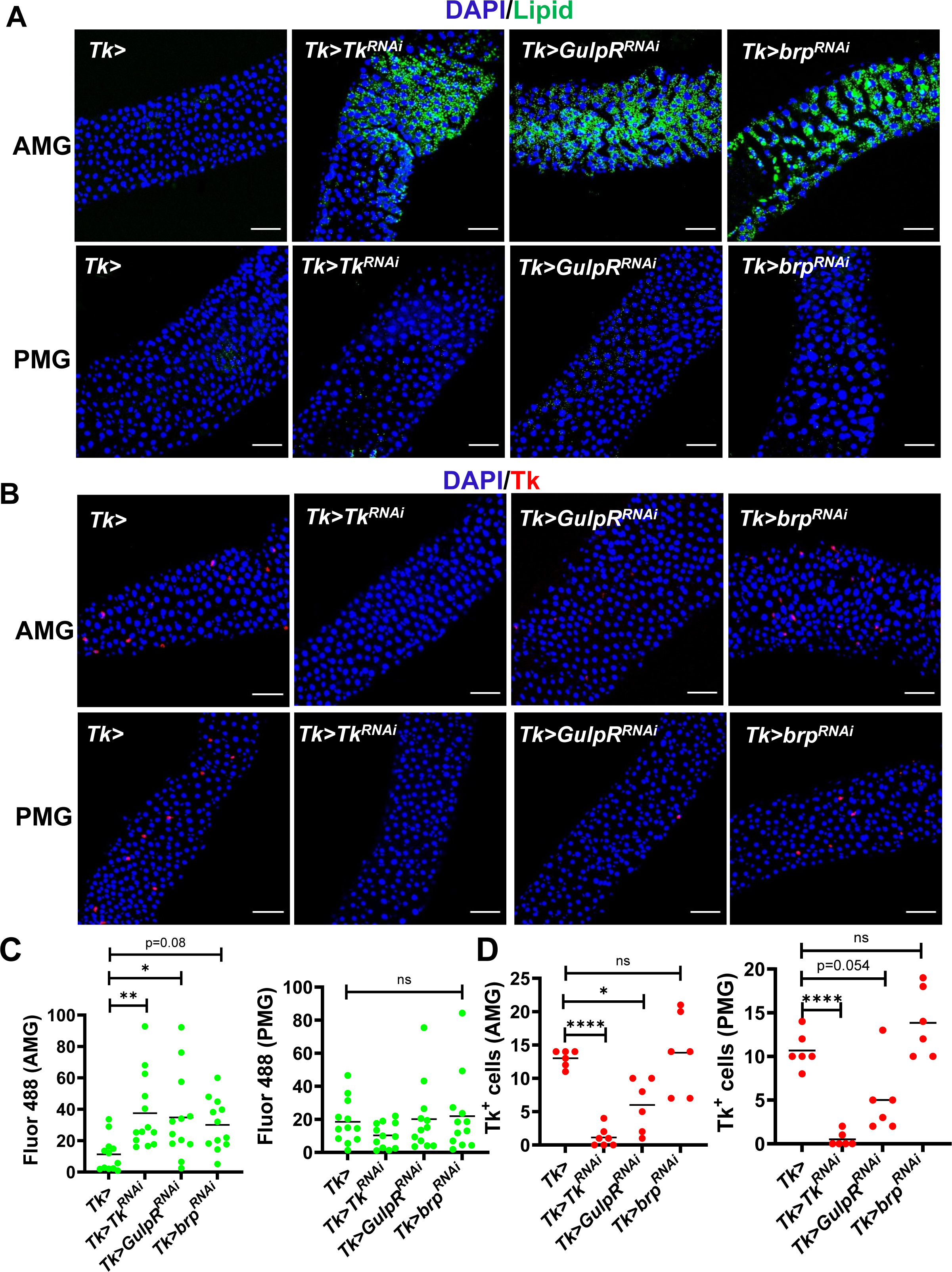
GulpR^RNAi^ in Tk+ EECs decreases Tk expression. (A) Representative fluorescence images showing DAPI and (A) BODIPY(Lipid) staining or (B) Tk immunofluorescence in the anterior midgut (AMG) and posterior midgut (PMG) of the indicated fly genotypes. Scale bars, 50 μM. Quantification of (C) total fluorescence and (D) Tk+ cells in the AMG and PMG of the indicated flies. The mean of at least six intestines is shown. For lipid quantification, a one-way ordinary ANOVA was used to assess significance. For Tk+ cells, a Welch’s ANOVA with a Dunnett’s T3 multiple comparisons test was used. **** p < 0.0001, ** p < 0.01, * p < 0.05, ns not significant.

Because this had not previously been reported, we questioned whether accumulation of lipids in the AMG only represented region-specific Tk regulation of lipid homeostasis or a direct action of GulpR [10]. To explore this, we compared lipid accumulation and Tk staining in the intestines of Tk>*Tk*^RNAi^ flies with that in Tk>*GulpR*^RNAi^ and Tk> intestines. In fact, Tk+ cell numbers were decreased in the intestines of Tk>*Tk*^RNAi^ flies beyond that observed for Tk>*GulpR*^RNAi^ flies but lipid accumulation was observed only in the AMG (Fig 2). These results indicate that GulpR in Tk+ cells regulates Tk expression and/or secretion and that Tk expression regulates lipid homeostasis in the AMG only. GPCRs may regulate EEP secretion as well as expression. Therefore, we examined the possibility that a block in secretion might result in negative feedback on expression leading to a decrease in Tk+ cells. Bruchpilot (brp) is an exocytosis factor that is essential for peptide secretion [21, 22]. To test the phenotype of a block in secretion, we compared the phenotype of Tk>*brp*^RNAi^ flies with that of Tk>*GulpR*^RNAi^ flies. Inhibition of Tk exocytosis via *brp*^RNAi^ resulted in lipid accumulation in the AMG but no change in the number of Tk+ EECs (Fig 2). This suggests that a block in Tk release does not result in decreased *Tk* expression and supports a role for *GulpR* in activating Tk expression rather than release.

### GulpR activates expression of additional EEPs produced by Tk+ EECs

To further differentiate regulation of EEP transcription and secretion, we conducted an RNA sequencing experiment comparing the intestinal transcriptomes of Tk>*GulpR^RNAi^* and Tk>*brp^RNAi^* flies to that of Tk> control flies. Using a threshold of 2-fold change and a padj of 0.05, 82 genes were differentially regulated by *GulpR*^RNAi^ with 41 increasing in transcription and 41 decreasing (Figure 3A and Table S1). Using the same criteria, *brp*^RNAi^ differentially regulated 124 genes with 78 increased and 46 decreased (Fig 3B and Table S2). Twenty-nine genes were similarly regulated by the two suggesting overlapping but distinct functions. Genes whose transcription was uniquely decreased in the intestines of Tk>*GulpR^RNAi^* flies included *Tk* as well as two additional EEPs expressed in the same EEC subtype: *Neuropeptide F* (*NPF*) and *Diuretic hormone 31* (*Dh31*). In contrast, none of these reached our threshold in Tk>*brp^RNAi^* flies. These results were confirmed by qRT-PCR (Fig 3C and D). We measured a 1.8-fold decrease in *Dh31* transcription in the intestines of Tk>*brp*^RNAi^ flies. A similar pattern was observed in the RNAseq experiment, suggesting that the action of brp or a product released from Tk+ cells regulates *DH31* transcription. To establish that Tk does not regulate transcription of *DH31* and *NPF* directly, we measured transcription of these genes in Tk> and Tk>*Tk*^RNAi^ flies. While *Tk*^RNAi^ did decrease transcription of *Tk* significantly, no change in transcription was observed for *DH31* and *NPF* (Fig 3E). We conclude that *Dh31* and *NPF* transcription is not regulated by Tk and that GulpR directly regulates EEP expression rather than secretion.

**Figure 3:**
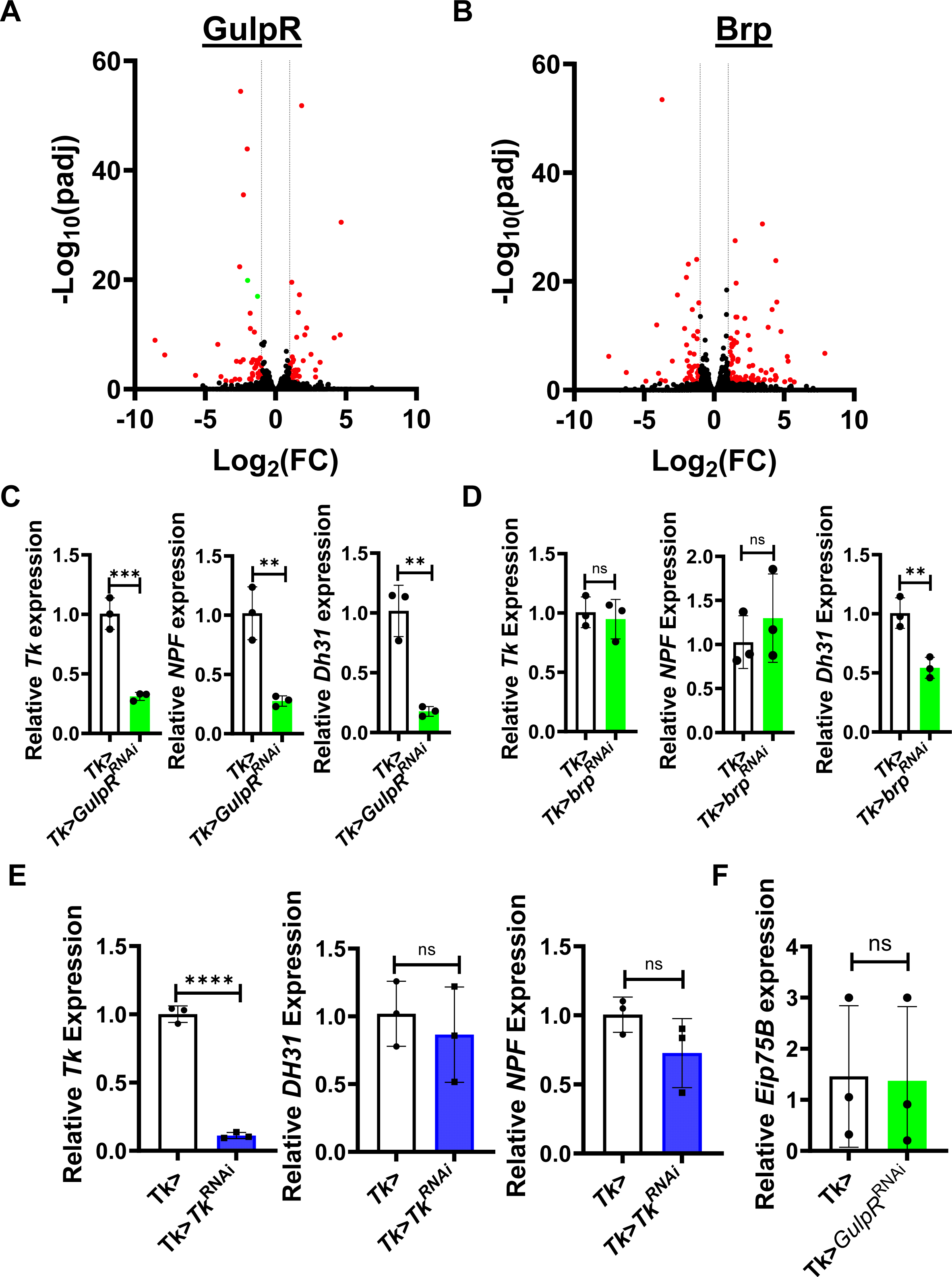
*GulpR*^RNAi^ in Tk+ EECs alters expression of innate immune genes and EEPs. Volcano plots of gene expression ratios derived from RNAseq analyses. (A) Tk>*GulpR*^RNAi^/Tk> or (B) Tk>*brp*^RNAi^/Tk>. Dotted lines indicate thresholds of 2-fold differences in transcription, while red dots represent genes that were significantly differentially regulated. (C-E) qRT-PCR analysis of genes encoding enteroendocrine peptides expressed by Tk+ cells in the indicated flies lines. (F) qRT-PCR analysis of the ecdysone-regulated gene *Eip75B* in the indicated fly lines. The mean of biological triplicates is shown. Error bars indicate the standard deviation. A student’s t test was used to assess significance. **** p < 0.0001, *** p < 0.001, ** p < 0.01, ns not significant.

We have previously uncovered signaling pathways that co-regulate IMD activation of *Tk* expression and ecdysone signaling [15, 23]. No ecdysone-regulated genes were identified in our RNAseq analysis. Nevertheless, to demonstrate that GulpR regulates *Tk* transcription independent of ecdysone signaling, we measured transcription of the ecdysone-regulated gene *Eip75B*, in Tk> and Tk>*GulpR*^RNAi^ flies. As shown in Fig 3F, *GulpR*^RNAi^ had no impact on *Eip75B* transcription. Therefore, we have identified a novel signaling pathway in control of EEP expression that is independent of ecdysone signaling.

### Intestinal GulpR and Tk are essential for regulation of lipid utilization during starvation

During long term starvation and infection, organisms use lipid stores to satisfy their energy requirements. We hypothesized that GulpR might play a role in one or both of these processes. To test this, we characterized Tk>, Tk>*GulpR*^RNAi^, and *Tk*^RNAi^ flies under nutrient-replete and nutrient-limited conditions. We found that total lipid stores were similar in Tk>, Tk>*GulpR*^RNAI^ and Tk>*Tk*^RNAi^ flies under nutrient-replete conditions (Fig 4A). As expected, lipid stores decreased in response to nutrient limitation (Fig 4B). However, we found that, under nutrient-limited conditions, lipid stores were significantly greater in Tk>*GulpR*^RNAi^ and *Tk*^RNAi^ flies as compared with control flies (Fig 4C). This suggests that both GulpR and Tk are essential for appropriate mobilization of lipid stores during starvation. We hypothesized that appropriate use of lipid stores during starvation might be essential for survival of nutrient stress. In fact, we found that Tk>*GulpR*^RNAI^ and Tk>*Tk*^RNAi^ flies succumbed to starvation more rapidly than Tk> control flies (Fig 4D).

**Fig 4:**
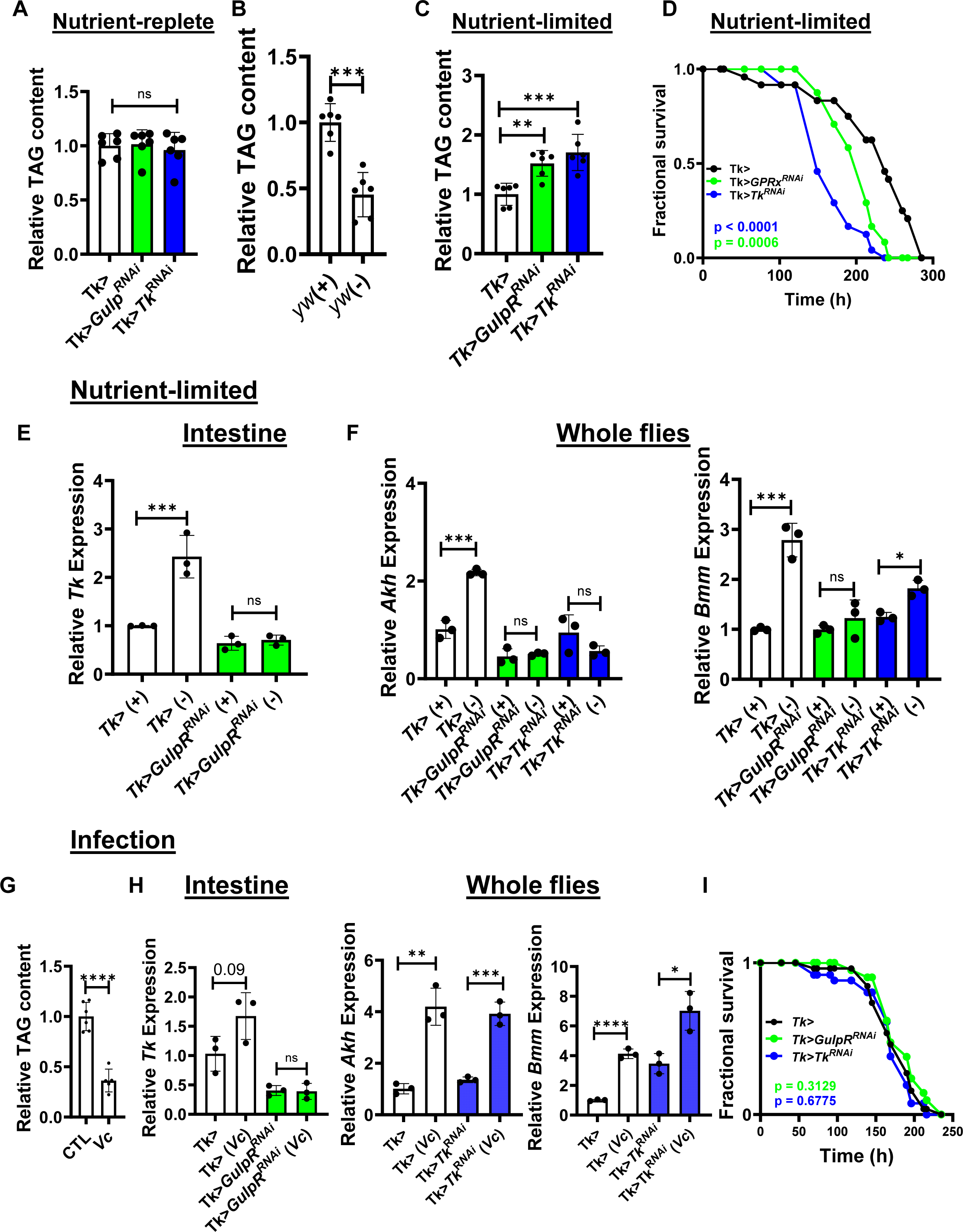
GPRx regulates lipid mobilization during starvation but not infection. Quantification of systemic triglycerides (TAG) in flies of the indicated genotypes after 72 hours of maintenance in (A) nutrient-replete, (B) nutrient-replete (+) and nutrient-limited (-), and (C) nutrient-limited conditions. The mean of at least five biological replicates is shown. Error bars represent the standard deviation. For A and C, an ordinary one-way ANOVA with Dunnett’s multiple comparisons test was used to assess significance. For B, a student’s t test was used. (D) Survival of nutrient-limitation over time for flies with the indicated genotypes. Thirty flies were tested per condition. Log-rank analysis was used to assess significance. (E) qRT-PCR analysis of transcription of the indicated genes in the intestines of flies with the indicated genotypes under fed (+) and nutrient-limited (-) conditions for 72 h. (F) qRT-PCR analysis of transcription of the indicated genes in whole flies with the indicated genotypes under fed (+) and nutrient-limited (-) conditions for 72 h. (G) Quantification of systemic triglycerides (TAG) in *yw* flies after 72 hours of infection with *V. cholerae* (*Vc*). The mean of at least five biological replicates is shown. Error bars represent the standard deviation. A student’s t test was used to assess significance. qRT-PCR analysis of transcription of the indicated genes in the intestines of flies with the indicated genotypes under uninfected or *V. cholerae* infected (*Vc*) conditions. (H) qRT-PCR analysis of transcription of the indicated genes in whole flies with the indicated genotypes under uninfected or infected (*Vc*) conditions. The mean of biological triplicates is shown. Error bars represent the standard deviation. A student’s t test was used to assess significance. (I) Survival of *V. cholerae* infection over time for flies with the indicated genotypes. Thirty flies were tested per condition. Log-rank analysis was used to assess significance. **** p<0.0001, *** p<0.001, ** p<0.01, * p<0.05, ns not significant.

Because GulpR regulates *Tk* transcription and both play a role in survival of starvation, we questioned whether GulpR might regulate *Tk* transcription in response to nutrient-limitation. To test this, we measured *Tk* transcription in the intestines of Tk> and Tk>*GulpR*^RNAi^ flies. As shown in Fig 4E, nutrient limitation significantly increased intestinal *Tk* transcription in Tk> flies, but this increase was not observed in Tk>*GulpR*^RNAi^ flies. The glucagon homolog Adipokinetic hormone (Akh) is responsible for mobilization of energy stores from the fat body in response to starvation, and Brummer (Bmm) is the main TAG lipase in the fly [24, 25]. We, therefore, hypothesized that GulpR and Tk might activate transcription of these two genes in response to nutrient limitation. To test this, we measured systemic transcription of *Akh* and *bmm* in Tk> control, Tk>*GulpR*^RNAi^, and Tk>*Tk*^RNAi^ under both nutrient-replete and nutrient-limited conditions. Nutrient limitation resulted in transcriptional activation of *Akh* and *Bmm* in Tk> control flies but not in Tk>Tk*^RNAi^* and Tk>*GulpR*^RNAi^ flies (4F). This suggests that GulpR directly regulates *Tk* under conditions of starvation resulting in activation of *Akh* and *Bmm* transcription to appropriately mobilize lipid stores.

### Intestinal GulpR and Tk are dispensable for lipid utilization during infection with the intestinal pathogen *Vibrio cholerae*

Pathogenic infection activates the host immune response in a process that is energetically costly [26, 27]. Similar to starvation, flies also lose significant amounts of their TAG stores during *V. cholerae* infection (Fig 4G). Having established that GulpR regulates intestinal *Tk* and that Tk regulates systemic *Akh* and *Bmm* during starvation, we reasoned that it might play a similar role during infection with the intestinal pathogen *Vibrio cholerae* [28]. While *V. cholerae* infection modestly increased transcription of *Tk* in the gut and this was blocked by Tk>*GulpR^RNAi^*, Tk did not control systemic *Akh* and *Bmm* transcription in the setting of infection (Fig 4H). Furthermore, neither GulpR nor Tk played a role in survival of infection (Fig 4I). We conclude that, while GulpR regulates *Tk* transcription in the setting of infection, increases in systemic *Akh* and *Bmm* transcription are regulated via a distinct pathway. As a result, GulpR plays no role in survival of infection.

## Discussion

EECs primarily function in expression and secretion of EEPs, which modulate local and systemic physiology. Regulation of EEP production relies on the ability of EECs to sense intestinal nutrient status, which is mainly achieved via nutrient transporters and GPCRs. In addition to nutrients, EEC GPCRs can sense and respond to other factors present in the intestine, including inflammatory cytokines, gut peptides, and neurotransmitters. Previously, we identified the EEC transporter Tarag, which functions as an acetate importer to activate expression of Tk [15]. Here, we characterize GulpR, an EEC GPCR that regulates expression of EEPs, including Tk, in the setting of metabolic stress.

Although we have not identified the ligand that activates GulpR signaling here, two studies suggest the type of ligands that might interact with GulpR. GulpR has been reported to be closely related to Neuropeptide Y receptors and prolactin-releasing peptide receptor [29]. Neuropeptide Y and prolactin-releasing peptide are both peptide hormones produced mainly in the vertebrate brain with some expression in the intestine [30–32]. In addition, a phylogenetic study suggested that GulpR was closely related to a *Drosophila* orphan GPCR CG12290, which is a predicted octopamine receptor [33]. We hypothesize that the endogenous ligand of GulpR is either a neuropeptide or a neurotransmitter.

EEPs are important regulators of metabolic homeostasis. There are two Tk receptors that are expressed in the gut, TkR99D and TkR86C. Tk interacts with its cognate receptor TkR99D on enterocytes to prevent accumulation of lipids in the intestine [10]. In this study, we found that during starvation the *Drosophila* EEP Tk is transcriptionally upregulated and essential for transcriptional activation of two main lipid utilization genes Akh and Bmm. One possibility is that Tk could be acting directly on the endocrine cells of the corpora cardiaca (CC), which are the main producers of Akh [34]. These cells express the Tk receptors TkR86C and TkR99D, and it has been reported that addition of Tk to the corpora cardiaca (CC) of *Locusta migratoria* ex vivo stimulates Akh release [35, 36].

Much like starvation, infection by the enteric pathogen *V. cholerae* induces GulpR-dependent upregulation of *Tk.* However, the significant increases in *Akh* and *Bmm* that are activated by infection are present even in the absence of Tk and neither Tk nor GulpR promotes survival of infection. Our work, therefore, suggests that there is redundancy in the signaling pathways that activate *Akh* and *Bmm* transcription to regulate lipid mobilization. However, while lipid mobilization is regulated mainly by the GulpR/Tk pathway in the setting of starvation, during infection, GulpR and Tk play minor roles and other signaling pathways dominate.

## Materials and Methods

### *Drosophila* Husbandry and Strains

Fly lines used in this study were fed standard Bloomington recipe fly food containing 16.5 g/L yeast, 9.5 g/L soy flour, 71 g/L cornmeal, 5.5 g/L agar, 5.5 g/L malt, 7.5% corn syrup, and 0.4% propionic acid. Flies were raised in a 12 hr day-night cycle incubator at 25°C. Adult female flies between 4-7 days of age were used in all experiments. The control lines *TRiP* (BL36303) and *yw* (BL1495) and the *GulpR* mutant stock (*GulpR^f06408^*:BL18976) were obtained from the Bloomington *Drosophila* Stock Center (BDSC). The following RNAi fly stocks were also obtained from the BDSC: *GulpR^RNAi^* (BL28621), *Tk^RNAi^* (BL2500), and *brp^RNAi^* (BL25891). The *Tk-*Gal4 driver line was a kind gift from Norbert Perrimon.

### Starvation-resistance

Three cohorts of 10 flies were placed and kept in vials containing an autoclaved cellulose plug infused with 3mL of 1x PBS. For survival, the number of viable flies was recorded twice daily until the number of viable flies was 0 for all genotypes. For RT-qPCR and TAG assays, the flies were kept in the vials containing 1x PBS for 72 hours and then processed for the appropriate experiment.

### V. cholerae infection

The quorum sensing-competent *V. cholerae* 01 El Tor strain C6706 (C6706 str 2) was used for fly infections. *V. cholerae* was grown in LB broth supplemented with 100 µg/mL streptomycin at 27°C. Three cohorts of 10 flies were orally infected with a 10-fold dilution of an overnight culture of *V. cholerae* in LB broth as follows. Flies were placed in vials containing an autoclaved plug infused with 3mL of the bacterial suspension and allowed to ingest it continuously. The number of viable flies was recorded twice daily.

### *Drosophila* RNA extraction, RNA-sequencing, and RT-qPCR

RNA was isolated from the intestines or whole bodies of 4-7 day old flies that were kept in standard fly food, starved for 72 hours by placing in vials containing cellulose plugs infused with 3 mL of 1x PBS, or infected with quorum sensing-competent *V. cholerae* 01 El Tor strain C6706 (C6706 str2) for 72 hours. Total RNA was isolated from 10-15 fly intestines or 5-8 whole flies per replicate using TRIzol reagent (Thermo Fisher Scientific 15596026) and the Direct-zol RNA Miniprep plus kit (Zymo Research R2070). A minimum of three biological replicates per condition/genotype were performed. For RNA sequencing analysis, the RNA was submitted to the Molecular Biology Core Facilities (MBCF) at the Dana Farber Cancer Institute (DFCI) for next-generation sequencing (NGS) library preparation, sequencing, and analysis (https://mbcf.dana-farber.org/totalrnaseq.html). Libraries were prepared using Roche Kapa mRNA HyperPrep strand specific sample preparation kits from 200ng of purified total RNA according to the manufacturer’s protocol on a Beckman Coulter Biomek i7. The finished dsDNA libraries were quantified by Qubit fluorometer and Agilent TapeStation 4200. Uniquely dual indexed libraries were pooled in an equimolar ratio and shallowly sequenced on an Illumina MiSeq to further evaluate library quality and pool balance. The final pool was sequenced on an Illumina NovaSeq X Plus targeting 40 million 150bp read pairs per library. Sequenced reads were aligned to the UCSC hg38 reference genome assembly and gene counts were quantified using STAR (v2.7.3a) [37] and Salmon [38]. Differential gene expression testing was performed by DESeq2 (v1.22.1) [39]. RNAseq analysis was performed using the VIPER snakemake pipeline [40].

For qPCR, cDNA was synthesized from 500ng of total RNA using the iScript™ cDNA synthesis kit (Bio-Rad 1708891). qPCR of target gene transcripts was performed using the iTaq™ Universal SYBR^®^ Green Supermix (Bio-rad 1725121) on either a StepOnePlus Real-Time PCR system (Applied Biosystems) or a QuantStudio™ 5 Real-Time PCR system (Thermo Fisher Scientific). Quantification cycle values (Cq) were obtained and used to calculate target gene transcription normalized to Actin. Primers used in this study are listed in Table S3.

### Immunofluorescence

Fly intestines were dissected in 1x PBS, fixed in 4% paraformaldehyde (PFA) solution (4% PFA in 1x PBS-0.1%Tween 20 (PBT)) for 20 minutes, and washed three times for 10 minutes with 1x PBT. For Tk-immunofluorescence experiments, intestines were left in blocking solution (PBT + 0.1% Triton X-100 (Sigma-Aldrich 9002-93-1) + 2% BSA (Sigma-Aldrich 9048-46-8)) for 1 hour, and then in Rabbit anti-Tk primary antibody solution (blocking solution + 1:500 anti-Tk antibody) overnight at 4°C. The next day the guts were washed three times with PBT for 10 minutes, followed by incubation in staining solution #1 (blocking solution + 1:1000 DAPI (Invitrogen D1306) + 1:500 Alexa 594-conjugated Goat anti-rabbit secondary antibody (Thermo Fisher Scientific A11012)) for 2 hours. For lipid staining, the intestines were incubated in staining solution #2 (PBT + 1:1000 DAPI (Invitrogen D1306) + 1:1000 BODIPY 493/503 (Invitrogen D3922)) for 2 hours after the initial fixing and washing steps. After three 10-minute washes in PBT, the guts were mounted in Vectashield® antifade mounting medium (Vector Laboratories H-1000-10) and imaged using a Zeiss LSM 980 confocal microscope and a 40x oil objective. Unless otherwise noted, all steps were done at room temperature. Tk+ cells were quantified manually. Lipid accumulation was assessed using ImageJ (FIJI) to quantify total BODIPY fluorescence. The corrected total cell fluorescence (CTCF) was calculated by dividing total BODIPY fluorescence by the total area and subtracting background fluorescence from this measurement. A minimum of six fly intestines per genotype were evaluated for quantification of Tk-expressing cells and BODIPY fluorescence.

### Triglyceride quantification

Where indicated, flies were starved or infected with *V. cholerae* for 72 hours prior to the measurement. Three cohorts of 5 flies were washed with cold 1x PBS in a 9-well glass plate. The flies were then homogenized in 100µL of cold 1x PBS-0.05% Tween using a plastic pellet pestle (Fisher Scientific) and a Fisherbrand™ Pellet Pestle™ Cordless Motor (Fisher Scientific). The homogenate was kept on ice until a heat-inactivation step involving incubation at 70 °C for 10 min. After heat-inactivation, 20µL of 1x PBS-0.05% Tween buffer or 20µL of triglyceride reagent (Sigma-Aldrich T2449) were added to 20µL of the homogenate and the mixture was incubated at 37°C for 1 hour. The samples were centrifuged at maximum speed for 3 min and 30µL of the supernatant were transferred to a clear-bottom 96-well plate. These samples were treated with 100µL of free glycerol reagent (Sigma-Aldrich F6428). Absorbance was measured at 540nm using a SpectraMax^®^ ABS absorbance microplate reader (Molecular Devices). Relative triglyceride (TAG) levels were calculated by subtracting the absorbance of samples treated with buffer (free glycerol) from samples treated with triglyceride reagent (free glycerol + fatty acids), dividing by the number of flies in each cohort (5), and normalizing to the control genotype.

### Quantification and statistical analysis

All data was graphed and analyzed using GraphPad Prism 10.0 software. Measurements shown in each graph represent the mean values of at least three biological replicates and the error bars represent ± the standard deviation. A Student’s t-test, a one-way ANOVA statistical test, or a Log-rank test was used to determine significance where appropriate. The statistical test used in each graph is specified in the figure legend.

### Data availability statement

RNAseq data have been deposited in the NCBI repository (accession no. GSE294931).

## Acknowledgements

This work was supported by NIH R01AI158247 and NIH R01AI162701 to P.I.W. and NIH F31DK130254 to D. B. Anti-TK antibodies were generously provided by Jan Veenstra. The TK-Gal4 and NP1-Gal4 (Myo1A-Gal4) driver flies were kind gifts from Norbert Perrimon. Stocks obtained from the Bloomington Drosophila Stock Center (NIH P40OD018537) were used in this study. Microscopy images were acquired at the Microscopy Resources on the North Quad (MicRoN) core at Harvard Medical School. We thank Paola Montero Lopis at the MicRoN core for providing expertise with image acquisition and quantification.

## Supplementary Materials

**Table S1: Genes differentially regulated in RNAseq of uninfected Tk>GulpR RNAI flies as compared with Tk> driver only flies (Tk>RNAi/Tk>). Driver-only flies were crossed to a Trip control fly line (y sc v).**

**Table S2: Genes differentially regulated in RNAseq of uninfected Tk>*brp*^RNAI^ flies as compared with Tk> driver only flies (Tk>RNAi/Tk>). Driver-only flies were crossed to a Trip control fly line (y sc v).**

**Table S3: Primers used in qRT-PCR experiments**

